# Synergistic Antitumor Activity of SUMO1 degrader and Oxaliplatin through G6PD Inhibition in Colorectal Cancer

**DOI:** 10.1101/2025.04.08.647644

**Authors:** Sunghan Jung, Rakhim Roy, Yin Quan Zhao, Thomas M. O’Connell, Regan K. Wohlford, Daeho Kim, Chunhai Hao, Anita C. Bellail

## Abstract

Colorectal cancer (CRC) stands as one of the primary causes of death despite advancements in targeted therapies and chemotherapeutic methods. The oxaliplatin-based chemotherapy, such as the FOLFOX, remains limited by its toxic side effects and the development of drug resistance. The current studies demonstrate redox homeostasis as an important therapeutic target in cancer cells through the discovery of glucose-6-phosphate dehydrogenase (G6PD) as their major controller of oxidative stress and survival. In this study, we investigated the therapeutic potential of HB007 in combination with FOLFOX in CRC. Treatment with HB007 decreased the enzyme activity of G6PD and generated high ROS concentrations, which subsequently induced intrinsic apoptosis. When combined with FOLFOX, which also induces ROS and inhibits G6PD activity, HB007 showed synergistic cytotoxicity in vitro, in colon patient-derived 3D organoids, and in vivo patient-derived xenograft model, including FOLFOX-resistant tumors. The combination treatment mechanistically targeted G6PD activity, disrupted redox balance, and activated apoptosis without affecting G6PD protein level. These findings suggest that G6PD inhibition by HB007 enhances the efficacy of FOLFOX, suggesting a strategy to overcome chemoresistance and improve therapy in advanced CRC.

## Introduction

Colorectal cancer (CRC) is still a leading cause of cancer mortality despite many achievements regarding the identification of potential therapeutic targets^1,2^. Of the standard treatments, FOLFOX, which involves the use of 5-fluorouracil, leucovorin, and oxaliplatin as key components, is widely used, especially in the late stages of the CRC^3^. Oxaliplatin is a platinum-based chemotherapeutic agent that triggers apoptosis of cancer cells^4^. It does that mainly through the production of ROS, which causes oxidative stress and the interference of essential cellular operations^5^. However, the application of oxaliplatin in clinical practice often meets problems of chemoresistance and significant side effects, including peripheral neuropathy, that restrain the long-term usage of oxaliplatin in patients^6–8^. Such issues provide a strong rationale for the development of new therapeutic approaches that would increase the potential benefits of chemotherapeutic treatments while minimizing toxicity.

Over the past few years, attention has been raised to the strategy that involves the incorporation of conventional chemotherapy in combination with molecules that have a selective affinity to specific molecular targets believed to influence tumor cell survival and resistance^9,10^. Among the pathways, the preservation of the redox balance in cancer cells has attracted a lot of interest, with glucose-6-phosphate dehydrogenase (G6PD) playing an important role in this process^11^. G6PD is an enzyme that belongs to the pentose phosphate pathway, and its function is critical as it provides NADPH, a molecule that is necessary for maintaining the redox state of the cell through the reduction of glutathione^12^. Due to elevated metabolic activity, cancer cells have high levels of oxidative stress^13,14^. G6PD plays a pivotal role in controlling oxidative stress, hence facilitating the survival of cancer cells even under adverse environmental conditions^11,14,15^. Consequently, G6PD is a potential target for cancer treatment.

In our previous study, we identified G6PD as a novel molecular target of anticancer small molecule HB007 ^16^. Given G6PD’s role in maintaining redox balance and supporting cancer cell survival, we hypothesized that HB007 in combination with oxaliplatin may have a synergistic effect to induce apoptosis of CRC cells by amplifying oxidative stress. Here, we investigated the therapeutic potential of the combination treatment of HB007 and oxaliplatin in both colon cancer cellular models and a patient-derived xenograft model.

## Result

### HB007 selectively degrades SUMO1-conjugated proteins to exert anticancer activity in colorectal cancer

Our previous research identified the small-molecule degrader HB007 as a compound that reduces SUMO1-conjugated proteins specifically ^16^. To further investigate the therapeutic potential in CRC, we determined the SUMO1 expression patterns and clinical relevance. The elevated expression of SUMO1 in colon cancer patients significantly correlated with reduced overall patient survival (Figure 1A). Immunohistochemistry (IHC) demonstrated a stage-dependent increase in SUMO1 protein levels in CRC patient tissue (Figure 1B). IHC score was lowest in normal and benign tissues and progressively increased from Stage 1 to stage IV CRC tumors (Figure 1C). These data suggested that SUMO1 is increased progression and can be a therapeutic target in advanced CRC.

**Figure 1.**
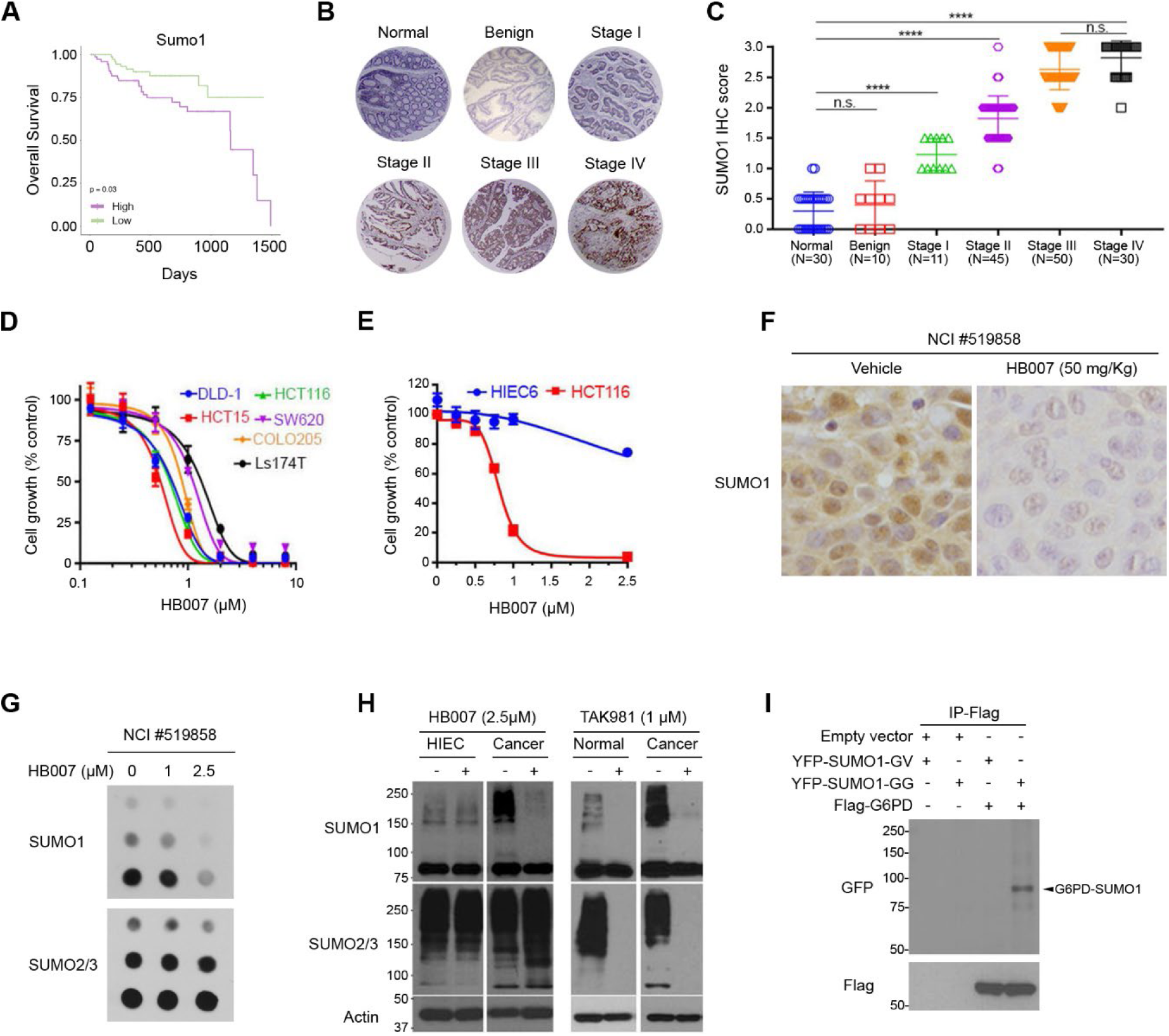
HB007 selectively targets and reduces SUMO1-conjugated proteins in colorectal cancer. (A) Kaplan–Meier survival analysis showing that high SUMO1 expression is associated with poor overall survival in CRC patients. (B) Representative IHC images of SUMO1 staining in human CRC tissues. (C) IHC scores of SUMO1 across CRC stages. SUMO1 expression progressively increased from normal and benign tissues to late-stage CRC (Stage II–IV). (D) Cell viability of HB007 in various CRC cell lines (DLD1, HCT15, SW620, COLO205, Ls174T, and HCT116). (E) HB007 selectively reduced viability in HCT116 cells while HIEC-6 showed lesser effect in dose-dependent manner. (F) IHC staining for SUMO1 in PDX tumors (NCI #519858) treated with vehicle or HB007 (50 mg/kg, intraperitoneal injection, 2 weeks). HB007 treatment markedly reduced SUMO1 staining intensity. (G) Dot blot analysis of PDX tumor lysates showing that HB007 selectively reduced SUMO1 but not SUMO2/3 levels in a dose-dependent manner. (H) Western blot analysis comparing the effects of HB007 (2.5 µM) and TAK981 (1 µM) on SUMO1 and SUMO2/3 levels in CRC cell and normal HIEC-6 cell. (I) Co-immunoprecipitation of Flag-G6PD with YFP-SUMO1 wild-type (GG) or conjugation-defective mutant (GV) in HCT116.

HB007 significantly decreased cancer cells growth in consistent with our previous data ^16^ and has lesser effect on normal intestinal epithelial cells (HIEC-6) (Figure 1D, 1E). In patient-derived xenograft model, IHC showed that HB007 administration reduced the SUMO1-ylated proteins in patient-derived xenograft (PDX) tumor (Figure 1F). Dot-blot assay of PDX tumors showed that HB007 reduced SUMO1, but not SUMO2/3 (Figure 1G). These results indicated that HB007 acts as a SUMO1 specific degrader in cancer.

We further examined specificity of HB007 compared to TAK981 (Subasumstat), known as SUMOylation inhibitors by inhibiting the SUMO-activating enzyme (SAE) ^17^. Western blot analysis showed that HB007 reduced the SUMO1-conjugated proteins in CRC cells selectively, while TAK981 decreased both SUMO1 and SUMO2/3-conjugated proteins in both CRC and normal cells (Figure 1H). These findings suggested that HB007 specifically targets and depletes SUMO1-conjugated proteins in CRC.

In our previous study, the genome-wide CRISPR-Cas9 knockout screens identified G6PD as top-ranked HB007-responsive proteins ^16^. To determine whether G6PD can be a target of HB007 as a SUMO1-ylated protein, co-immunoprecipitation assay was conducted and showed that G6PD was SUMO1-ylated (Figure 1I), as shown in a previous paper^18^. Given the identification of G6PD as a SUMO1-ylated protein and HB007-responsive protein, we next investigated its mechanistic roles in mediating the anticancer effect in CRC.

### HB007 inhibits G6PD and induces mitochondrial dysfunction via ROS accumulation in colorectal cancer

G6PD is a cytosolic enzyme that catalyzes the pentose phosphate pathway’s first and rate-limiting step ^15^. G6PD’s primary function is to produce nicotinamide adenine dinucleotide phosphate (NADPH), a key antioxidant for detoxification of high levels of reactive oxygen species (ROS) ^19,20^. Furthermore, G6PD activity is increased in several types of cancers, and elevated gene expression is commonly observed in many cancer types ^14^. To investigate the role of G6PD, we first analyzed the Cancer Genome Atlas (TCGA) and Genotype Tissue Expression (GTEx) database for G6PD expression in various human cancer types. G6PD mRNA expression was significantly increased in 16 of 31 cancers: breast invasive carcinoma (BRCA), cholangiocarcinoma (CHOL), colon adenocarcinoma (COAD), esophageal carcinoma (ESCA), glioblastoma multiforme (GBM), head and neck squamous cell carcinoma (HNSC), kidney chromophobe (KICH), Kidney renal papillary cell carcinoma (KIRP), lower grade glioma (LGG), liver hepatocellular carcinoma (LIHC), lung adenocarcinoma (LUAD), pancreatic adenocarcinoma (PAAD), rectum adenocarcinoma (READ), skin cutaneous melanoma (SKCM), stomach adenocarcinoma (STAD), and lung squamous cell carcinoma (LUSC) (Figure 2A). The significant increase of G6PD mRNA was further confirmed in an analysis of a large number of colon cancer and normal tissues (Figure 2B). This elevated expression of G6PD in colon cancer patients correlated significantly with reduced overall patient survival (Figure 2C). To further determine G6PD protein expression, western blot analysis of human colon cancer tissues and matched normal adjacent colon cancer tissues revealed elevated levels of G6PD protein in human colon cancer tissues (Figure 2D).

**Figure 2.**
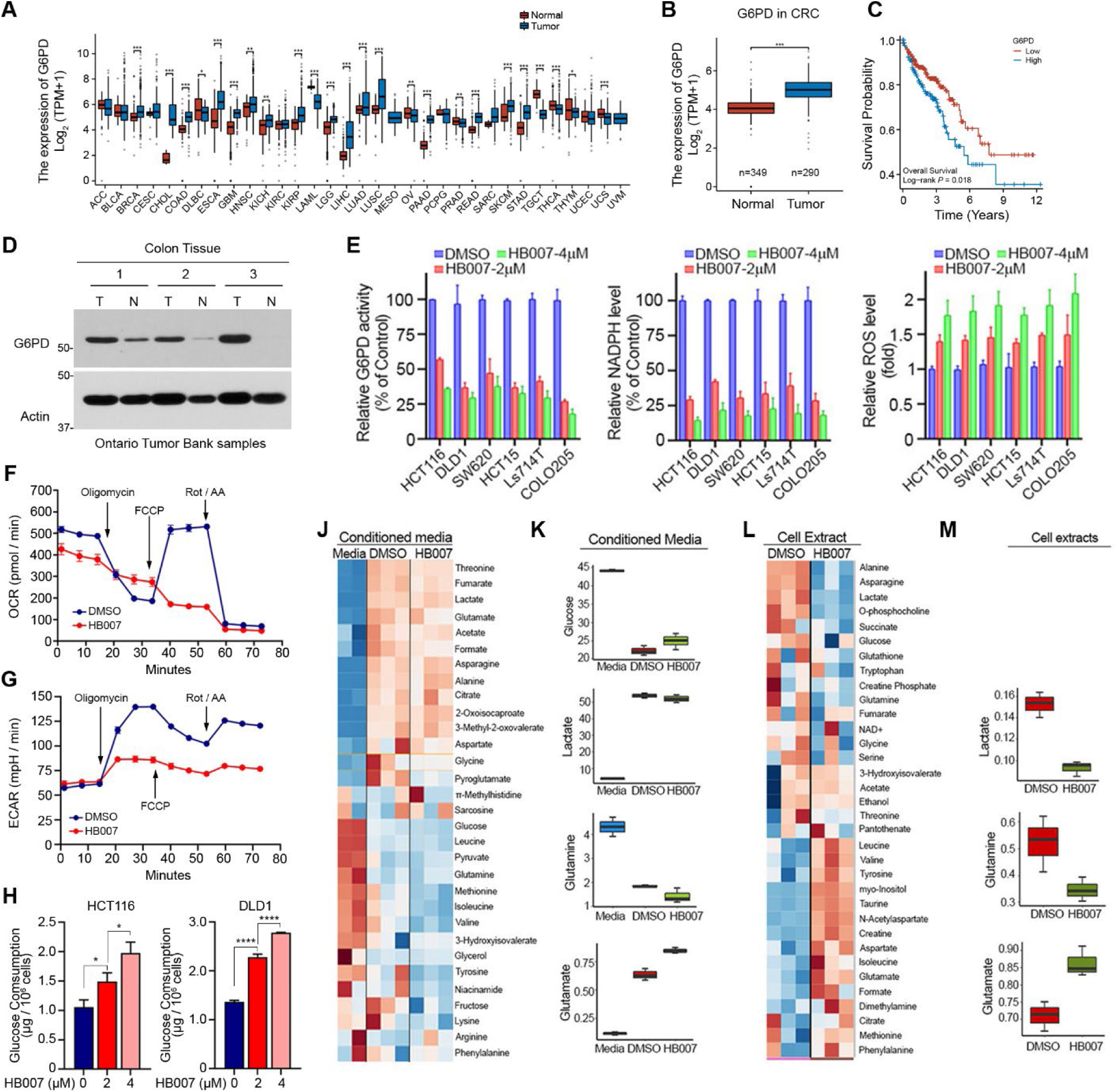
HB007 inhibits G6PD activity and induces mitochondrial dysfunction through ROS accumulation in colorectal cancer. (A–C) G6PD mRNA expression is significantly elevated in multiple cancers based on TCGA and GTEx datasets (A), with a marked increase in colorectal cancer (B) and associated with poor overall survival (C). (D) Western blot analysis shows increased G6PD protein levels in human colon tumors compared to matched normal tissues. (E) HB007 treatment significantly reduced G6PD activity and NADPH levels while increasing ROS in CRC cell lines (n=3). (F, G) Seahorse analysis revealed that HB007 reduces oxygen consumption rate (OCR) (F) and increases extracellular acidification rate (ECAR) (G) in HCT116 cells, indicating mitochondrial dysfunction. (H) HB007 increased glucose uptake in HCT116 and DLD1 cells in a dose-dependent manner (n=3). (J–M) NMR-based metabolomics analysis showed altered extracellular (J, K) and intracellular (L, M) metabolite profiles following HB007 treatment, including reduced lactate and glutamine levels. Data are presented as box plots or heatmaps.

To further determine the anticancer effects of HB007 through G6PD regulation, we treated HB007 on colon cancer cell lines. The results showed that HB007 significantly reduced G6PD activity and NADPH levels, while HB007 increased ROS levels in a dose-dependent manner in different colon cancer cell lines (Figure 2E).

G6PD produces NADPH to maintain redox equilibrium and supports mitochondrial metabolism^14,21,22^ and HB007 reduced NADPH and increased ROS in colon cancer cells (Figure 2E). To determine whether impaired redox buffering and oxidative damage may affect mitochondrial respiration, HCT116 was treated with HB007, and oxygen consumption rate (OCR) and glycolytic extracellular acidification rate (ECAR) were analyzed by Seahorse. The results showed that HB007 reduced OCR and indicated impaired mitochondrial respiration (Figure 2F). In contrast, ECAR increased suggested a compensatory upregulation of glycolytic flux in response to mitochondrial dysfunction (Figure 2G).

To confirm whether HB007 induces glycolytic compensation, colon cancer cells were treated with HB007, and glucose consumption was determined. HB007 led to an increase in glucose uptake in both HCT116 and DLD1 cells (Figure 2H). These results suggested that G6PD inhibition disrupts mitochondrial function and forces CRC cells to rely more heavily on glycolysis as an adaptive response.

To further investigate the metabolic impact of HB007 since G6PD is the rate-limiting step in the oxidative pentose phosphate pathway^15^, both conditioned media and intracellular extracts were analyzed by NMR (Nuclear Magnetic Resonance) based metabolomics analysis. Several significantly altered metabolites suggest specific modulation of cellular energetics. The extent of glucose uptake in the media increased after exposure to HB007 because of elevated glycolytic activity. The results from metabolomic profiling showed that lactate amounts inside the cells decreased significantly. Lactic acid is known to be accumulated in many cancer cells, including HCT116, and has been implicated in chemotherapeutic resistance and reduced apoptosis ^23^. Thus, a beneficial impact of HB007 treatment seems to be indicated by reduced intracellular lactate levels. The intracellular concentrations of glutamine lowering together with glutamate raising indicate that modifications to glutaminolysis likely contribute to the tumor-inhibiting properties of HB007. These indicated that cellular metabolism became impaired during HB007 treatment as part of its antitumor mechanism (Figure 2J-M). Together, these results demonstrated that HB007 impairs mitochondrial respiration through G6PD-related ROS and induces a shift toward glycolysis, accompanied by broad metabolic disruptions.

### Synergistic inhibition of G6PD by HB007 and oxaliplatin enhances anticancer activity in CRC cells

Since HB007 disrupts redox and metabolic homeostasis through G6PD inhibition, we next investigated whether G6PD could be targeted by combining HB007 with standard chemotherapeutic agents for colon cancers. Oxaliplatin is a critical component of the first-line standard chemotherapy regimen for colorectal cancer^24^. Oxaliplatin is known to induce DNA damage and mitochondrial ROS accumulation, and some studies have reported its potential influence on G6PD activity^25,26^. To determine whether oxaliplatin affects G6PD activity, we treated HCT116 with oxaliplatin and measured enzymatic activity. The results showed that oxaliplatin reduced G6PD activity in dose dose-dependent manner (Figure 3A), as well as HB007 (Figure 3B). Interestingly, Western blot analysis showed that HB007 and oxaliplatin did not change the total G6PD protein level in HCT116 (Figure 3C).

**Figure 3.**
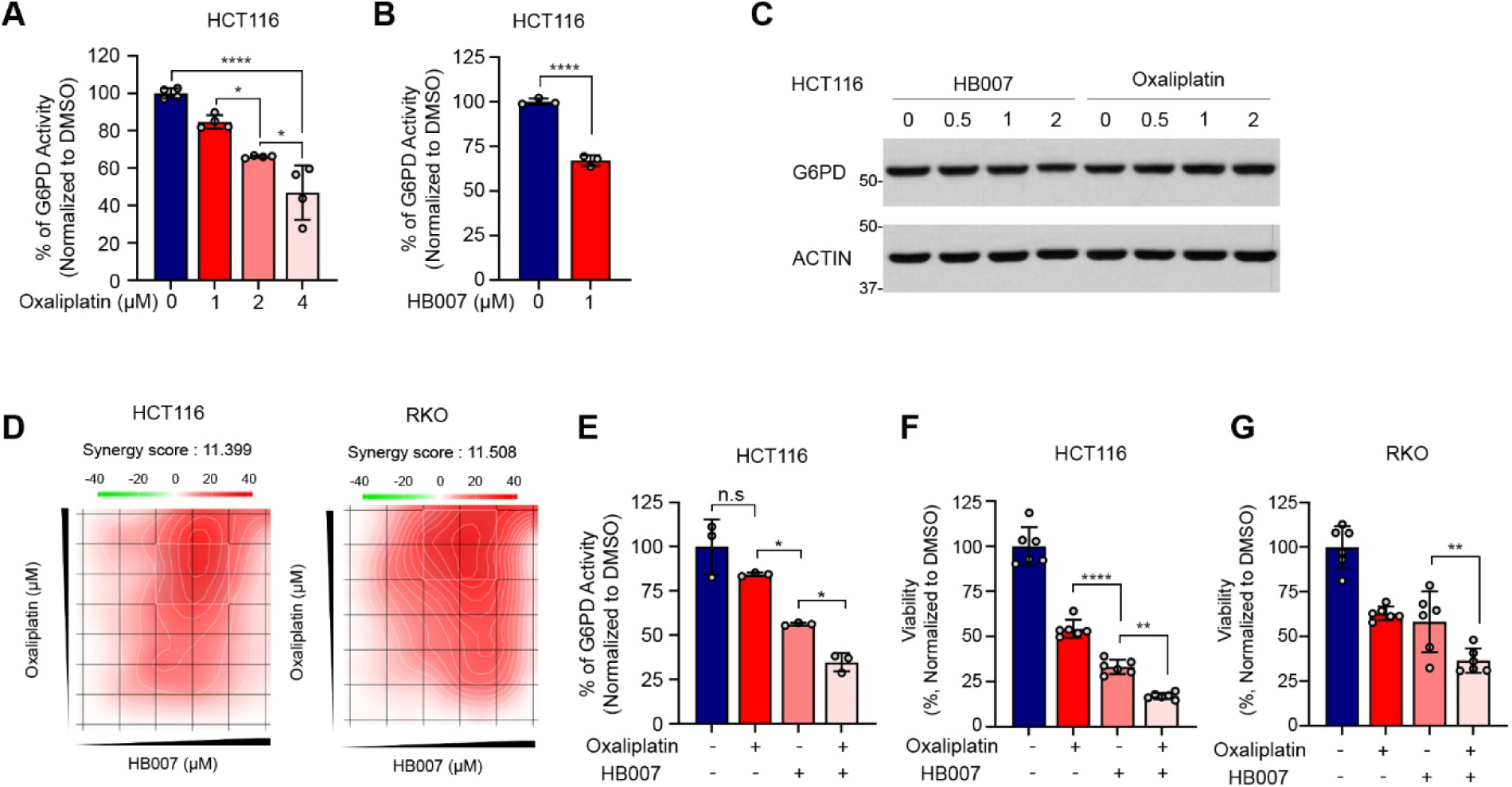
HB007 synergizes with oxaliplatin to inhibit G6PD and reduce CRC cell viability. (A–B) G6PD enzymatic activity in HCT116 cells decreased dose-dependently with oxaliplatin (A) and HB007 (B) (Indicated concentration for 2 days). (C) Western blot showing no change in total G6PD protein after treatment with HB007 or oxaliplatin (Concentration: indicated concentration for 2 days). (D) Synergy map and scores from combination treatment with HB007 and oxaliplatin in HCT116 and RKO cells using HSA model. (E) G6PD enzymatic activity was significantly lower in the combination group than in single treatments (n = 3, HB007: 1 µM, oxaliplatin: 2 µM for 2 days). (F–G) Cell viability significantly decreased with HB007 and oxaliplatin combination compared to single treatments in HCT116 and RKO (n = 6, HB007: 1 µM, oxaliplatin: 2 µM for 2 days).

A previous study reported that suppression of G6PD expression enhances the sensitivity of colon cancer cells to oxaliplatin^26^. Since our data showed that both HB007 and oxaliplatin reduced G6PD activity in single treatments, we hypothesized that co-treatment of HB007 and oxaliplatin would show a synergistic effect on G6PD activity, reducing cell viability in colon cancer cells. We treated HB007 and oxaliplatin in several combination doses to HCT116 and RKO and analyzed the cell viability value to get the synergy score, choosing the best combination treatments and showing an 11.399 synergy score and 11.0508 in HSA analysis methods, respectively (Figure 3D). G6PD activity was reduced more in combination treatments compared to either single treatment in HCT116 (Figure 3E). To determine the functional consequence of this combination on G6PD activity, we carried out a cell viability assay in HCT116 and RKO cells. The combination treatment significantly reduced cell viability compared to single treatment in both HCT116 and RKO (Figure 2F, 2G). These results suggested that oxaliplatin contributed to G6PD activity inhibition and enhanced the anticancer effect of HB007 through enhancing ROS targeting in colon cancer cells.

### Co-treatment with HB007 and oxaliplatin promotes ROS accumulation and intrinsic apoptosis in colorectal cancer cells

Since both HB007 and oxaliplatin reduced G6PD activities and enhanced cell death in our previous data, we next examined the mechanisms underlying this synergistic cytotoxicity in colorectal cancer cells. Since G6PD is critical for regulating redox balance through NADPH production^14,26,27^, we hypothesized that HB007 and oxaliplatin combination treatment would result in excessive ROS accumulation, mitochondrial damage, and intrinsic apoptotic cell death. To determine this hypothesis, first, ROS levels were measured in HCT116. Both HB007 and oxaliplatin increased ROS levels in a dose-dependent manner in 1-day treatment and 1-hour treatments (Figure 4A, 4B). Co-treatment results further amplified ROS accumulation (Figure 4C). It suggested a synergistic disruption of redox homeostasis with combination treatments.

**Figure 4.**
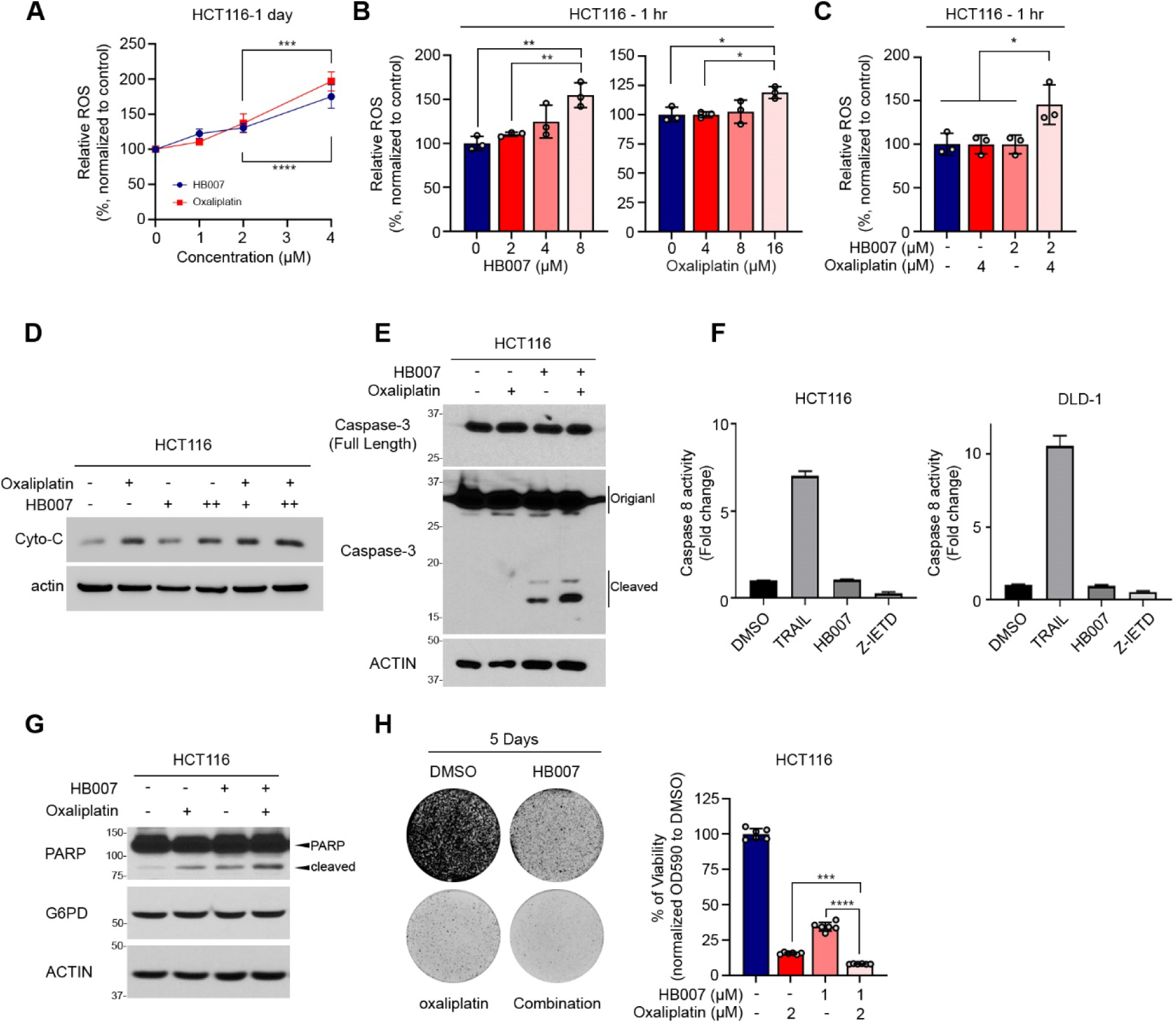
HB007 and oxaliplatin co-treatment increases ROS and induces intrinsic apoptosis in CRC cells. (A–C) ROS levels measured by DCF-DA assay increased dose-dependently with HB007 and oxaliplatin; co-treatment further amplified ROS in HCT116 cells (n = 3). (D) Western blot showing increased cytosolic cytochrome c after combination treatment (HB007: 1 µM and 2 µM, oxaliplatin: 2 µM for 2 days). (E) Cleaved caspase-3 increased upon co-treatment, confirming the intrinsic apoptosis pathway. (F) Caspase-8 activity assay showed no significant change with HB007, indicating no involvement of extrinsic apoptosis (n = 3). TRAIL and Z-IETD were used as positive and negative controls, selectively. (G) Western blot analysis showed that cleaved PARP increased with combination treatment, without altering G6PD protein levels (HB007: 1 µM, oxaliplatin: 2 µM for 2 days). (H) Colony formation assay showed near-complete abrogation of colony growth with combination treatment in HCT116. Bar graph quantifies percent viability.

Furthermore, an excessive ROS can damage mitochondria and trigger intrinsic apoptotic pathways by releasing cytochrome C from mitochondria into the cytosol ^28,29^. Therefore, we determined the released cytochrome c levels by lysing cells with mild buffer, followed by western blot analysis. The combination treatment increased cytosolic cytochrome C levels compared to single treatments or control (Figure 4D). This indicated that combination treatment induced mitochondrial outer membrane permeabilization and promoted activation of the intrinsic apoptotic pathway.

To confirm the apoptotic pathway, we analyzed cleaved caspase-3 (an executor caspase activated during apoptosis) and cleaved PARP (poly ADP-ribose polymerase; a downstream substrate cleaved by active caspase-3). The results showed that the combination treatment increased cleaved caspase-3 in HCT116 (Figure 4E). Oxaliplatin was reported to activate caspase-8 in a previous paper^30^. To determine whether HB007 may also affect on extrinsic apoptotic pathway, we conducted a Caspase-8 activity assay, but HB007 showed no significant change in colon cancer cells (Figure 4F). Without changing the amounts of G6PD protein, combination treatments increased PARP cleavage activity (Figure 4G). The results of the crystal violet assay demonstrated that the combination treatment induced drastic cell death and blocked colony formation (Figure 4H). These findings suggested a mechanism by which HB007 and oxaliplatin cooperatively target G6PD to impair NADPH production and ROS regulation. This disruption led to the accumulation of intracellular ROS, mitochondrial dysfunction, and the intrinsic apoptotic pathway, inducing cell death.

### Genetic suppression of G6PD sensitizes colorectal cancer cells to HB007 and oxaliplatin via ROS-mediated apoptosis

Since our findings showed G6PD is a key regulator for the synergistic anticancer effect of compounds, we next investigated the functional role of G6PD using genetically modified cell models. To confirm whether G6PD is necessary for the antitumor effect of HB007 and its synergy with oxaliplatin, we generated G6PD-deficient HCT116 cells using shRNA-mediated knockdown. Western blot analysis showed that the G6PD protein level decreased in shG6PD HCT116 clones compared to shControl HCT116(Figure 5A). shG6PD HCT116 showed efficiently suppressed G6PD activity in two different clones (Figure 5B). Cell growth assay showed that G6PD knockdown significantly reduced cell proliferation in two different clones (Figure 5C), consistent with the findings in G6PD knockdown in other cancer cells ^26,31,32^. ROS levels were elevated in shG6PD HCT116 clones (Figure 5D). Next, to confirm whether increased ROS derived from G6PD knockdown induces more apoptotic cell death, we treated HB007 and oxaliplatin to shControl and shG6PD HCT116 in single and combination treatments, followed by western blot analysis of cleaved PARP. The results showed that shG6PD cells increased PARP cleavages and further increased with compound treatments (Figure 5E). These findings indicated that loss of G6PD activity sensitizes the compound’s anticancer effect. To verify this, we used CRISPR-Cas9-mediated G6PD knockout HCT116 (Figure 5F). We found similar results with PARP cleavage analysis in G6PD knockout HCT116 cells treated with HB007 along with oxaliplatin when compared to shControl HCT116 cells (Figure 5G). These data indicate that G6PD loss increases sensitivity to oxidative stress and induces apoptosis, which supports that G6PD is the molecular target for HB007 and oxaliplatin cytotoxicity.

**Figure 5.**
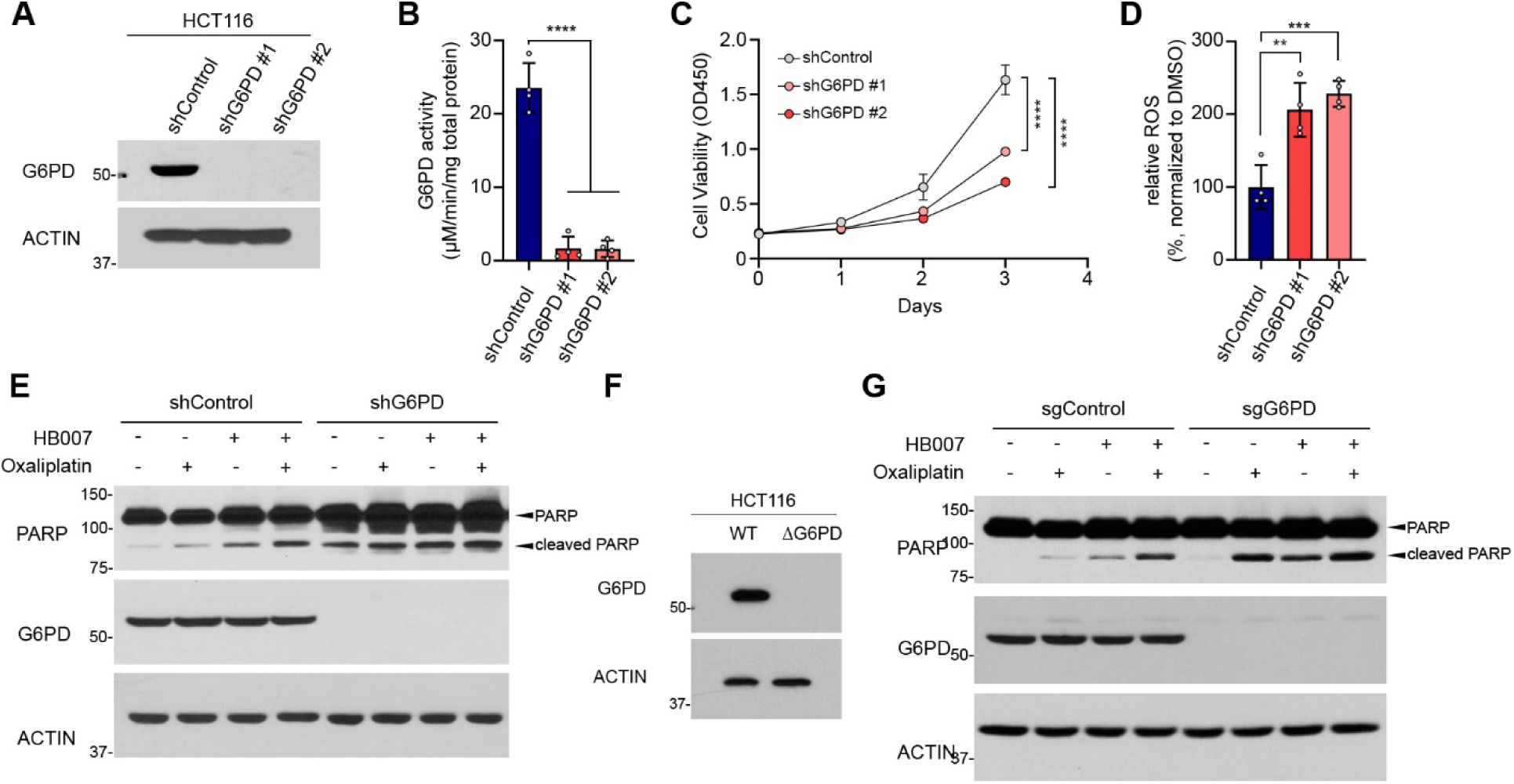
Genetic suppression of G6PD sensitizes CRC cells to HB007 and oxaliplatin-induced apoptosis. (A–B) Western blot and G6PD activity assay confirm effective G6PD knockdown in HCT116 via shRNA (n = 3). (C) Cell proliferation was significantly reduced in shG6PD clones over 3 days (n = 3). (D) ROS levels were significantly increased in shG6PD clones compared to control (n = 3). (E) Western blot analysis showed that Cleaved PARP levels were increased in shG6PD cells and further increased by HB007 or oxaliplatin treatment. (HB007: 1 µM, oxaliplatin: 2 µM for 2 days) (F) CRISPR-Cas9-mediated knockout of G6PD (ΔG6PD) in HCT116 cells showed a complete loss of G6PD protein expression, as confirmed by Western blot analysis. (G) In ΔG6PD HCT116 cells, treatment with HB007 and/or oxaliplatin led to increased cleaved PARP levels, with the combination treatment inducing the most significant apoptotic response (HB007: 1 µM, oxaliplatin: 2 µM for 2 days).

### HB007 synergizes with FOLFOX to inhibit G6PD and induce intrinsic apoptosis in patient-derived CRC 3D organoids

Building on our findings that the combination treatment of HB007 and oxaliplatin increases ROS accumulation and intrinsic apoptosis of colorectal cancer cells through a G6PD inhibition, we next investigated whether this occurred in human patient-derived colorectal cancer (CRC) 3D organoids^33^. CRC-3D organoids are a good model to demonstrate similarities to actual tumor structures and biological diversity ^34,35^. Following the well-established protocol^36,37^, we generated liver metastasized, lymph metastasized, and primary CRC-3D. The CRC-organoids were frozen, sectioned, and stained with transmembrane staining reagents, showing conventional 3D organoids morphology (Figure 6A). Liver and lymph metastasized 3D organoids were treated with HB007 and oxaliplatin in a dose-dependent manner. The viability results showed that CRC-3D organoids were decreased dose-dependently (Figure 6B, 6C).

**Figure 6.**
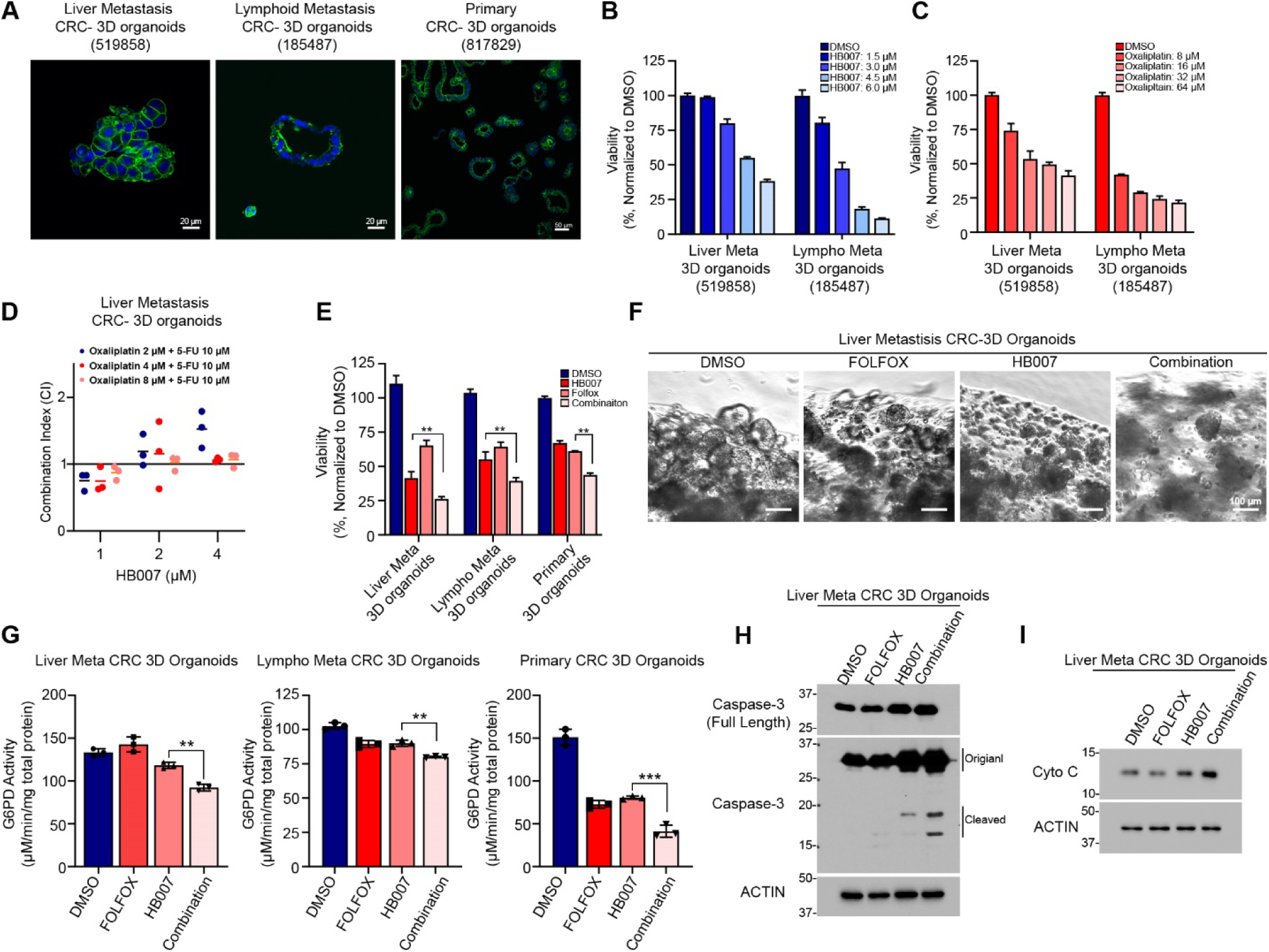
HB007 synergizes with FOLFOX to inhibit G6PD and induce apoptosis in CRC patient-derived 3D organoids. (A) Representative immunofluorescence images of CRC-3D organoids from liver metastasis, lymph node metastasis, and primary tumor. (B–C) Dose-dependent viability reduction in liver and lymph metastasis CRC-3D organoids after HB007 (B) and oxaliplatin (C) treatment (n = 3, treated for 3 days). (D) Combination index (CI) analysis confirms synergy of HB007 with FOLFOX components (treated in indicated doses for 3 days). (E) Viability assay showing enhanced cytotoxicity with triple combination (HB007 + oxaliplatin + 5-FU) across all organoid models (n = 3; HB007: 1 µM, oxaliplatin: 2 µM, 5-FU: 10 µM for 3 days). (F) Brightfield images showing enhanced organoid disruption in the combination group (HB007: 1 µM, oxaliplatin: 2 µM, 5-FU: 10 µM for 5 days). (G) G6PD enzymatic activity significantly reduced in all CRC-3D organoids after combination treatment (n = 3; HB007: 1 µM, oxaliplatin: 2 µM, 5-FU: 10 µM for 3 days). (H–I) Increased cleaved caspase-3 and cytosolic cytochrome c were detected by western blot in liver metastasis organoids (HB007: 1 µM, oxaliplatin: 2 µM, 5-FU: 10 µM for 3 days).

To further explore the translational relevance of HB007 in the context of standard clinical regimens, 5-fluorouracil (5-FU) was incorporated, which commonly serves as a crucial component of FOLFOX chemotherapy used for colorectal cancer treatment ^38,39^. First, a range of concentration combinations was tested, and the strongest synergistic effect was observed with HB007 (1 μM), oxaliplatin (2 μM), and 5-FU (10 μM), as indicated by combination index analysis (Figure 6D). This optimized combination showed significantly enhanced cytotoxicity across all three organoid models compared to single treatment (Figure 6E). Morphological assessment by brightfield microscopy further confirmed enhanced organoid disruption with the combination compared to single treatments (Figure 6F).

Next, to determine whether drug-induced G6PD inhibition observed in colorectal cancer cell lines also occurs in 3D organoid models, the activity of G6PD was evaluated in different types of CRC-3D organoid models following treatment with the combination of HB007 and FOLFOX. Consistent with earlier findings, the results showed the G6PD activities were significantly reduced after combining HB007 with FOLFOX treatment in three CRC-3D organoid models (Figure 6G).

To further determine whether the reduced G6PD activity induces intrinsic apoptotic cell death in CRC-3D organoids, as previously observed in colorectal cancer cells, we analyzed the cleaved caspase-3 and cytochrome C in western blot analysis. The results showed that cleaved caspase-3 levels and cytochrome C levels were markedly increased in the combination group compared to single treatments (Figure 6H). These data confirmed that HB007 with FOLFOX disrupts redox balance and mitochondrial integrity through G6PD inhibition in 3D tumor models derived from CRC patients.

### HB007 synergizes with FOLFOX to suppress tumor growth and improve survival in PDX models of colorectal cancer

In our previous studies, HB007 demonstrated a significant antitumor activity, significantly suppressing tumor growth in CRC-PDX models of colorectal cancer ^16,40^. To determine the combination treatment of HB007 and FOLFOX, 3 different types of CRC-PDX models (NCI #519858, NCI #185487, and NCI #997537) were employed. Drug treatment history showed that NCI #519858 and NCI #185487 experienced FOLFOX, but NCI #997537 were treatment naive. First, we administered the compounds via intraperitoneal injection (IP) for 2 weeks and monitored their tumor size, survival rate, and body weight assessments during 6-8 weeks. The combination treatment of HB007 and FOLFOX showed significant tumor regression compared to single treatments in all CRC-PDX models (Figure 7A, 7D, 7G). It also showed better survival rates (Figure 7B, 7E, 7H). During the treatment, no significant body weight changes were observed in the control group and the oxaliplatin group. Early treatment time in HB007 and the combination treatment group showed a slight body weight decrease, but it was restored during treatment (Figure 7C, 7F, 7I). These results demonstrated that the combination treatment has a synergic antitumor effect on PDX models.

**Figure 7.**
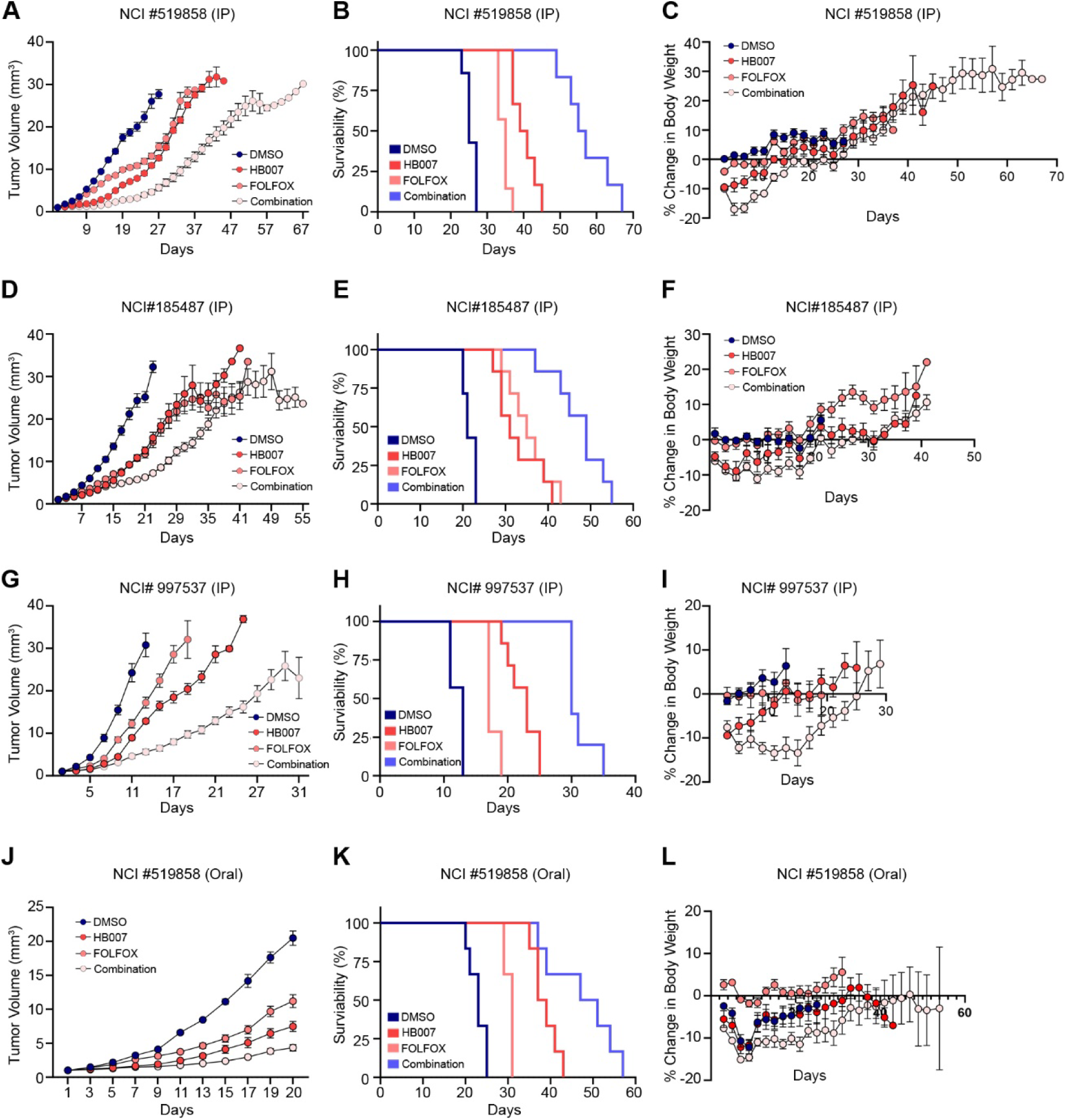
Combination of HB007 and FOLFOX suppresses tumor growth and improves survival in CRC PDX models. (A–C) NCI #519858 PDX model was treated with compounds, intraperitoneal (IP) co-treatment with HB007 and FOLFOX reduced tumor volume (A), improved survival (B), and showed minimal impact on body weight (C). (D–F) NCI #185487 PDX model was treated with compounds. Intraperitoneal (IP) co-treatment with HB007 and FOLFOX reduced tumor volume (D), improved survival (E), and showed minimal impact on body weight (F). (G–I) In treatment-naïve model NCI #997537, combination treatment led to significant tumor suppression and survival benefit (G–H) with tolerable weight change (I). (J–L) Oral administration of HB007 combined with FOLFOX in NCI #519858 also reduced tumor burden (J), extended survival (K), and had manageable body weight effects (L). All in vivo experiments used n = 6–8 mice per group. Statistical analysis was performed using two-way ANOVA or Kaplan–Meier survival analyses.

Next, PDX-model (NCI #519858) was administered by oral injection and was monitored. The combination treatment showed a significant tumor regression compared to the single and control, and improved survival rate (Figure 7J, 7K). Oral formulation reduced bodyweight early, but it was slightly restored during treatment (Figure 7L). Together, these data highlight the broad and robust antitumor activity of HB007 in vivo and support its potential as a combination partner with standard chemotherapy in both treatment-resistant and naïve colorectal cancer models.

## Discussion

According to our findings, HB007-mediated inhibition of G6PD increases the sensitivity of CRC cancer cells to oxidative stress and enhances the efficacy of oxaliplatin-based chemotherapy. HB007 disrupts redox homeostasis by suppressing G6PD activity, inducing ROS accumulation, and intrinsic apoptotic cell death in CRC cells. Oxaliplatin, known to induce ROS ^41,42^, also reduces G6PD activity in our results. And when it is used in combination with HB007, it shows synergistic cytotoxic effects. This synergy is likely derived from dual suppression of the antioxidant system. This combination robustly disrupts redox homeostasis and triggers intrinsic apoptotic cell death than single treatment. These findings were further supported by shRNA and CRISPR-mediated G6PD loss-of-function cell models. It exhibited more sensitivity to both compounds. This effect occurred without changes in total G6PD protein level, but with G6PD enzymatic activity changes.

These findings are further strengthened by CRC patient-derived 3D organoids, which show more accurate recapitulated structure, heterogeneity, and drug response of human tumors ^33,35^. It also showed synergistic cytotoxic effect in combination therapy in organoids derived from both primary and metastatic CRC tissues. Consistent with cancer cell line results, CRC-3D organoids also show synergistic antitumor effect with HB007 and FOLFOX. This combination treatment also showed its synergistic efficacy in the FOLFOX-resistant PDX model. It suggests that HB007 may help overcome acquired chemoresistance by targeting metabolic vulnerabilities. Furthermore, in vivo studies showed synergistic antitumor effects with the combination of HB007 and FOLFOX.

Several limitations should be mentioned despite these findings. Our data demonstrate the importance of G6PD as a key target, however, the direct binding of HB007 to G6PD, as well as more molecular targets of HB007, remains to be fully elucidated. While we verified redox imbalance and intrinsic apoptosis, further investigation into additional signaling pathways (AMPK^43,44^ and NRF2^45,46^, and mitochondrial biogenesis^47,48^) must be necessary because it would lead to a more comprehensive mechanistic understanding. Long-term efficacy and the potential development of resistance to HB007-based therapies may be essential for assessing its clinical durability.

This research shows that G6PD inhibitor HB007 with oxaliplatin creates a new treatment method that disrupts redox balance to break through CRC chemoresistance. In vitro and in vivo experiments have shown robust evidence that HB007 presents great prospects as an effective combined agent for treating progressive colorectal cancer.

## Materials and Methods

### Cell lines and reagents

Human colorectal cancer cell lines HCT116, RKO, DLD1, SW620, Ls714T, COLO205, and HIEC-6 were obtained from ATCC and cultured in DMEM or RPMI-1640 medium supplemented with 10% FBS and 1% penicillin/streptomycin. Oxaliplatin was purchased from Selleck Chemicals (Houston, TX) and dissolved in water, and HB007 was dissolved in DMSO.

### RNA expression analysis

transcriptome data from the TCGA and GTEx datasets were analyzed using the XIANTAO web platform (www.xiantao.love). Briefly, RNA-seq data were obtained from the UCSC XENA browser and processed using the Toil pipeline^49^. TPM values were log2-transformed as log2(TPM + 1), and paired comparisons between cancer and adjacent normal tissues were performed across multiple cancer types to compare G6PD gene expression.

### G6PD knockdown HCT116 generation

Stable knockdown of G6PD was generated using lentiviral shRNA constructs targeting human G6PD (Sigma). Transduced HCT116 cells were selected with puromycin (2 µg/mL).

### G6PD activity test

G6PD activity was measured using the G6PD Activity Assay Kit (Abcam) according to the manufacturer’s protocol or Fang et al protocol ^50^. In brief, the lysis buffer contained HEPES 20 mM (pH 7.8), NaCl 150 mM, 0.5% Triton X-100, and glycerol 5%. Each well received lysate amounts between 5 and 7 µg protein with 180 microliters of activation buffer containing 50 mM Tris (pH 7.5), 5 mM MgCl₂, 1 mM glucose-6-phosphate (G6P), and 0.1 mM NADP⁺. It was measured by using a Synergy H1 microplate reader (BioTek) at 340 nm excitation with 460 nm emission every 5 minutes through 30–60 minutes.

### DCF-DA assay (ROS detection)

Cells were seeded into a black/clear bottom 96-well plate. Compounds were treated in phenol red-free media for 24 h. Cells were incubated with 25 μM DCF-DA diluted in PBS for 45 min at 37°C in CO_2_ incubator. DCF-DA solution was removed, and cells were washed by PBS once. The signal from cells was detected by a microplate reader at excitation 485 nm/emission 535 nm.

### Seahorse metabolic flux assay

Mitochondrial respiration and glycolysis were evaluated using a Seahorse XF Analyzer (XF96, Agilent). 10K/well cells were seeded a day ahead. Oxygen consumption rate (OCR) and extracellular acidification rate (ECAR) were measured following HB007 treatment (2 µM). For OCR, oligomycin (1 µM), FCCP (2 µM), and rotenone/antimycin A (1 µM) were sequentially injected.

### Metabolomics

NMR-based metabolomic analysis was performed on conditioned media and intracellular extracts of HCT116 cells treated with HB007. Samples for NMR experiments were prepared using an established protocol^51^. NMR spectra were acquired using a Bruker AVANCE III 700 MHz spectrometer equipped with a cryogenically cooled probe. Metabolite concentrations were normalized to total protein content.

### Cell viability and colony formation assays

Cell viability was determined by CCK-8 assay and phosphatase assay^40^. Cells were seeded into 96-well plates and treated with compounds for indicated time. Absorbance was read at 450 nm and 405 nm, selectively. For long-term survival, cells were seeded in 6-well plates, treated with HB007 and/or oxaliplatin, cultured for 5 days, fixed with methanol, and stained with 0.5% crystal violet (25% MeOH in DDW).

### Western blot analysis

Cells or tissues were lysed in RIPA buffer containing protease inhibitor cocktails and phosphatase inhibitors (Sigma #3). Samples were resolved by using SDS-PAGE and transferred to NC membranes. Membranes were probed with antibodies against G6PD (Proteintech, CST), PARP (CST), cytochrome C (BD Biosciences), and β-actin (CST).

### Caspase-8 activity assays

Caspase-8 activity followed the protocol supplied by the Caspase-8 Fluorometric Assay Kit (BioVision). Briefly, cells were lysed in the lysis buffer in the kit and incubated on ice for 10 min. Equal amounts of proteins were added to a reaction buffer containing10 mM DTT and IETD-AFC substrate. Then, the mixture was incubated at 37°C for 1 to 2 hours in the dark. The fluorescence was measured using a microplate reader (440 nm/505 nm).

### 3D organoid generation, culture, western blotting

PDX tissues were harvested from mice and minced into tiny pieces. Minced tissues were incubated at 37°C for 30 min to 1 hr with tissue dissociation solution (Collagenase II 1.5 mg/ml, Hyaluronidase 20 μg/ml, and DNase I 200U / HBSS with Ca 2+, Mg 2+). Dissociation buffer was inactivated by 3D organoid washing media (Advanced DMEM/F12 Medium with HEPES (1M) GlutaMAX (200 mM), 1% Penicillin/Streptomycin, 2.5% Fetal Bovine Serum (heat inactivated)) and passed through a 100 μm cell strainer. Dissociated cells were washed with PBS and plated with Matrigel embedding. After Matrigel dorms became solid, 3D organoid culture media (Advanced DMEM/F12 Medium with HEPES (1M) GlutaMAX (200 mM), 1% Penicillin/Streptomycin, B-27 supplement, 10% R-spondin 1 conditioned media, 125 μM N-acetylcysteine, 5 ng/ml EGF, 0.1 μg/ml Noggin) were added in well plates. For western blotting of 3D organoids, 3D organoids were harvested and extracted from Matrigel. Proteins were extracted by RIPA with sonication or mild lysis buffer (0.5% Triton X-100) with 20 min incubation on ice.

### ICC for 3D organoid

3D organoids were harvested from cell culturing plates and extracted from organoid-bearing Matrigel. Organoids were fixed in 10% formalin at 4°C overnight. After the fixation, organoids were permeabilized by 30% sucrose solution (in PBS) at 4°C overnight, embedded in optimal cutting temperature compound (OCT), and then the samples were kept in –80°C. The frozen samples were sectioned using a cryostat and collected on ultra-frosted glass microscope slides. Sectioned samples were permeabilized by 0.3% Triton-X100 in PBS, blocked by 0.1% Triton-X100 PBS solution with 10% v/v BSA. Primary antibodies were incubated in 0.1% Triton-X100 PBS solution with 5% v/v BSA; subsequently, Alexa Fluor-conjugated secondary antibodies (goat anti-rabbit or mouse IgG) in PBS were used to target the specific proteins.

### Mice experiment

Mice experiments were approved by the Institutional Animal Care and Use Committee of the Indiana University School of Medicine. HB007 and FOLFOX (5FU, oxaliplatin) were dissolved in 25% DMSO, 25% Kolliphor EL, and 50% PBS for intraperitoneal injection. For oral administration, HB007 was diluted in 5% 1-Methyl-2-pyrrolidinone, 10% propylene glycol, 5% Kolliphor R-HS15, 10% sulfobutylether-β-cyclodextrin (SBECD, Captisol) in water (5:10:5:80). PDXs fragments were obtained from NCI Patient-Derived Models Repository and tissues were implanted subcutaneously in right flanks of NOD/SCID gamma (NSG) mice (Jackson). When the implanted tissues reached 50 mm^3^ to 70 mm^3^, mice were allocated into groups and treated with HB007 or vehicle once per day for 14 days or treated once a week for 2 times for FOLFOX (5-FU 40 mg/kg; oxaliplatin 2.4 mg/kg) following the previous protocol^52^. The size of the tumor was measured by digital calipers every 2 days, and the volume of the tumor was calculated by (length × (width)^2^)/2. For the survival test, mice were checked until the tumor volume reached 1500 mm^3^. Mice were monitored for 6 to 8 weeks after the implantation. For Oral administration, compounds were dissolved in 1-methyl-2-pyrrolidinone (NMP), propylene glycol (PG), Kolliphor® HS-15 (Solutol), 10% sulfobutylether-β-cyclodextrin (SBECD, Captisol®) in water at a final % (10:10:5:75). Mice were administered the final solution via oral gavage at a 10 mg/kg concentration.

### Immunohistochemistry (IHC)

IHC was performed in the Department of Pathology and laboratory Medicine of Indiana University. Xenograft samples were fixed, paraffin-embedded, sectioned, and stained with SUMO1 antibody (Santa cruz).

### Statistical analysis

All statistical analyses were performed using GraphPad Prism 10 (GraphPad Software, La Jolla, CA). Data from in vitro experiments are presented as mean ± standard deviation (SD), while data from in vivo studies, including patient-derived xenograft (PDX) models, are presented as mean ± standard error of the mean (SEM). Comparisons between two groups were conducted using unpaired two-tailed Student’s *t*-tests. For comparisons among multiple groups, one-way or two-way analysis of variance (ANOVA) was used, followed by Tukey’s or Bonferroni’s post hoc test, as appropriate. Synergy scores for drug combinations were calculated using the Highest Single Agent (HSA) model in SynergyFinder^53^ or using the Chou-Talalay method (combination index methods) in compuSyn^54^. Kaplan–Meier survival curves were analyzed using the log-rank (Mantel-Cox) test. Statistical significance was defined as *p\** < 0.05; p** < 0.01; p*** < 0.001; p**** < 0.0001.

